# Biased signaling downstream of epidermal growth factor receptor regulates proliferative versus apoptotic response to ligand

**DOI:** 10.1101/352526

**Authors:** Remah Ali, Wells Brown, Stephen Conner Purdy, V. Jo Davisson, Michael K. Wendt

## Abstract

Inhibition of EGFR signaling by small molecule kinase inhibitors and monocloncal antibodies has proven effective in the treatment of multiple cancers. In contrast, metastatic breast cancers (BC) derived from EGFR-expressing mammary tumors are inherently resistant to EGFR-targeted therapies. Mechanisms that contribute to this inherent resistance remain poorly defined. Here we show that in contrast to primary tumors, ligand-mediated activation of EGFR in metastatic BC is dominated by STAT1 signaling. This change in downstream signaling leads to apoptosis and growth inhibition in response to EGF in metastatic BC cells. Mechanistically, these changes in downstream signaling result from an increase in the internalized pool of EGFR in metastatic cells, increasing physical access to the nuclear pool of STAT1. Along these lines, an EGFR mutant that is defective in endocytosis is unable to elicit STAT1 phosphorylation and apoptosis. Additionally, inhibition of endosomal signaling using an EGFR inhibitor linked to a nuclear localization signal specifically prevents EGF-induced STAT1 phosphorylation and cell death, without affecting EGFR:ERK1/2 signaling. Pharmacologic blockade of ERK1/2 signaling through the use of the allosteric MEK1/2 inhibitor, trametinib, dramatically biases downstream EGFR signaling toward a STAT1 dominated event, resulting in enhanced EGF-induced apoptosis in metastatic BC cells. Importantly, combined administration of trametinib and EGF also facilitated an apoptotic switch in EGFR-transformed primary tumor cells, but not normal mammary epithelial cells. These studies reveal a fundamental distinction for EGFR function in metastatic BC. Furthermore, the data demonstrate that pharmacological biasing of EGFR signaling toward STAT1 activation is capable of revealing the apoptotic function of this critical pathway.

## Introduction

Breast cancer metastasis is a multi-step process that culminates in vital organ invasion and proliferation by cancer cells. These later events of metastasis are responsible for patient morbidity and mortality in breast cancer ^1^. Developing targeted therapies for metastatic breast cancer faces many challenges. Paramount to these challenges is the high degree of molecular changes that characterize metastatic lesions compared to primary tumors, which constantly brings into question the utility of primary tumor analysis to guide metastatic therapy ^2,3^. Thus, understanding signaling events specific to metastatic breast tumors is essential to identify potential therapeutic targets and biomarkers for late stage disease.

Similar to more established breast cancer-associated genes, such as estrogen receptor (ER) and human epidermal growth factor receptor 2 (Her2), primary versus metastatic tumor discrepancies have also been described for epidermal growth factor receptor (EGFR)-expressing mammary tumors ^3–6^. Breast cancer cells predominantly respond to EGFR agonists in a proliferative fashion supporting its role as an oncogene. Indeed, studies from our group and others have linked activation of EGFR to mammary epithelial cell transformation, increased proliferation, and several early steps of metastasis ^7,8^. Various signaling pathways facilitate these oncogenic roles of EGFR, including the p38 mitogen activated protein kinase, extracellular signal-regulated kinases 1 and 2 (ERK1/2), signal transducer and activator of transcription 3 (STAT3), and phosphoinositide 3-kinase (PI3K). These experimental findings are supported by clinical studies that report high expression of EGFR in primary mammary tumors is predictive for reduced patient survival ^9,10^.

However, subsets of cancer cells, including those originating from the breast, respond to epidermal growth factor (EGF) via cell cycle arrest and induction of apoptosis ^11–14^ These observations are corroborated by the antitumor response of *in vivo* administered EGF ^15^. Many studies describe the growth-inhibitory functions of EGFR to be mediated by STAT1, which is an established tumor suppressor and mediator of apoptosis downstream of interferon signaling ^16–18^. We have recently shown that EGFR function changes from oncogenic in primary tumors to growth-inhibitory and apoptotic in metastatic tumors ^5,19^ The importance of this paradoxical function of EGFR is substantiated by the failure of EGFR inhibition (EGFRi) to improve the clinical outcomes of metastatic breast cancer patients ^20–28^. Inhibition of specific pathways downstream of EGFR is also being pursued for clinical applications. In particular, the compound trametinib is an allosteric inhibitor of MEK1/2, the kinases directly upstream of ERK1/2 ^29^. As opposed to direct inhibition of growth factor receptors, targeting of downstream pathways requires consideration that the cellular effects of inhibition may also arise via differential activation of alternate signaling pathways downstream of a common driver receptor.

In the current study, we demonstrate the apoptotic function of EGFR in metastatic breast cancer is dependent on STAT1 and we address the hypothesis that pharmacologic inhibition of MEK1/2 with trametinib will bias EGFR signaling toward a STAT1-dominated, apoptotic signaling pathway. These findings identify unique molecular signaling events that specifically manifest in metastatic BC, and identify a pharmacological approach to enhance STAT1-induced apoptosis and limit primary and metastatic tumor growth.

## Methods and Materials

### Cell lines and Reagents

Murine NMuMG and human MDA-MB-468 cells were purchased from ATCC and cultured in DMEM containing 10% or 5% FBS, respectively. MDA-MB-468 passages 1-5 were used in this study. NMuMG cells and their metastatic variants also received 10 μg/ml of insulin. Construction of NMuMG cells expressing human mutant (EGFR-AA) or WT-EGFR (NME) and their metastatic variants are described elsewhere ^7,30^. Cellular depletion of STAT1 cells was accomplished by VSVG lentiviral transduction of pLKO.1 shRNA vectors (Thermo Scientific), sequences of shRNAs can be found in Supplementary Table 1. The human MCF10-Ca1a cell line was kindly provided by Dr. Fred Miller (Wayne State University) and cultured in DMEM supplemented with 10% FBS. A list of the chemical inhibitors and growth factors used throughout the study can be found in Supplementary Table 2.

### Immunoblot and immunofluorescent analyses

For immunoblot assays, equal aliquots of total cellular protein were resolved by SDS-PAGE and transferred to PVDF membranes using standard methods. Immunofluorescent assays were conducted using primary antibodies in combination with fluorescently labeled secondary antibodies. Confocal images were captured using an Nikon A1Rsi inverted microscope. Super resolution images are structure illuminations obtained on a Nikon Ti-E inverted microscope with N-SIM capability. Antibody concentrations and suppliers are listed in Supplementary Table 3.

### Apoptosis and Cell viability assays

Caspase 3/7 activity was quantified using the Caspase-Glo 3/7 assay (Promega) according to manufactures instructions. Cell viability was measured using the CellTiter-Glo assay (Promega) according to manufactures instructions.

### 3D culture methods

Primary and metastatic breast cancer cells (5×10^3^) were plated into non-adherent roundbottom 96-well plates (Corning) in full growth media and cultured for 3 days. At this point the tumorspheres were physically transferred with 50 μl of residual media and 150 μl of fresh media to a flat bottom 96-well plate coated with 50 μl of growth factor reduced basement membrane hydrogel (Trevigen) in the presence or absence of trametinib and EGF. These structures were allowed to grow for an addition 3 days for metastatic NM-LM1 cells or 10 days for nonmetastatic NME cells at which point structure size was quantified using image J analyses or invasive tips were enumerated.

### Statistical methods

Statistical analyses were carried out using unpaired Student’s T-test or ANOVA where the data fit the parameters of the test. P values < 0.05 were considered statistically significant. P values for all experiments are indicated. All P values were generated using Prizm-Graph Pad.

## Results

### EGF-mediated STAT1 phosphorylation increases with metastasis

Our previous studies demonstrate that EGFR overexpression is capable of transforming normal murine mammary gland (NME) cells ^13,31_33^. Transient induction of epithelial-mesenchymal-transition (EMT) via treatment with TGF-β facilitates the metastasis and inherent resistance of these cells to EGFRi (fig. 1a; ^13^). To investigate differential downstream signals generated by EGFR in these isogeneic cells of increasing metastatic capacity, we examined the phosphorylation of STAT1 and ERK1/2 in response to exogenous EGF stimulation. As shown in Figure 1b and 1c, EGF treatment resulted in enhanced phosphorylation of STAT1 in lung-metastatic, LM1 and LM2, cells as compared to the nonmetastatic NME cells. In contrast, phosphorylation of ERK1/2 in response to EGF was similar in all cell types, therefore the ratio of ERK1/2 to STAT1 phosphorylation is dramatically altered in metastatic cells (fig. 1b and 1c). Importantly, the enhanced STAT1 signaling in the LM cells occurs despite EGFR returning to levels similar to endogenous in nontransformed NMuMG cells (fig. 1b). Unlike EGF, interleukin 6 (IL6) induced STAT3 activation was similar across all cells of the NME series (data not shown). Taken together, these findings suggest that through metastasis there is not a general propensity to increase STAT activation, but there is a specific increase in the EGFR:STAT1 signaling axis.

**Fig. 1.**
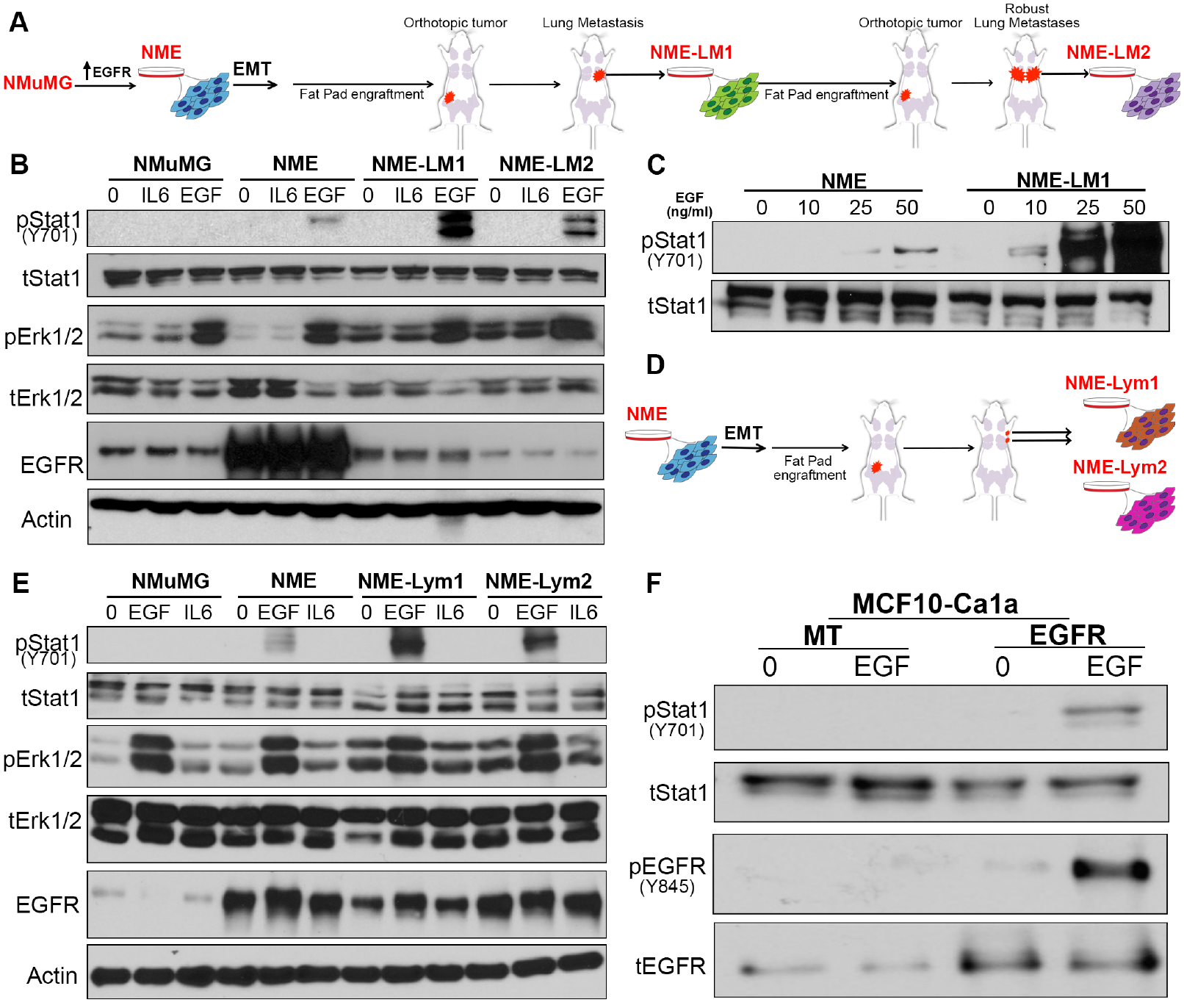
The STAT1:ERK1/2 signaling ratio downstream of EGFR is altered in metastatic cells compared to primary breast cancer cells. (**A**) Schematic representation of the EGFR transformed lung-metastatic breast cancer progression series. (**B**) Cells described in panel A were analyzed by immunoblot for phosphorylation of STAT1 and ERK1/2 in response to a 30 minute EGF stimulation (50 ng/ml). IL6 and BSA (0) served as protein stimulation controls, and analysis of total levels of EGFR, STAT1, ERK1/2 and Actin served as loading controls. (**C**) Non-metastatic (NME) and lung metastatic (NME-LM1) cells as described in panel A were stimulated for 30 minutes with the indicated concentrations of EGF and analyzed for phosphorylation of STAT1. Analysis of total STAT1 served as a loading control. (**D**) Schematic representation of the EGFR-transformed lymph node-metastatic breast cancer progression series. Metastatic cells were isolated from two separate lymph-nodes and termed NME-Lym1 and NME-Lym2. (**E**) Cells described in panel D were analyzed for phosphorylation of STAT1 and ERK1/2 in response to a 30 minute EGF stimulation (50 ng/ml). IL6 and BSA served as stimulation controls, and analysis of total levels of EGFR, STAT1, ERK1/2 and Actin served as loading controls. (**F**) EGFR was ectopically expressed in metastatic MCF10-Ca1a cells. These cells were stimulated with EGF (50 ng/ml) for 30 minutes and analyzed for phosphorylation of STAT1 and EGFR. Analysis of total levels of EGFR and STAT1 served as loading controls. All immunoblots shown are representative of at least three independent experiments.

To expand these observations, we derived additional metastatic lines from different anatomical locations. Metastases from two different lymph nodes were subcultured and termed NME-Lym1 and NME-Lym2 cell lines (fig. 1d). Consistent with our previously reported observations in lung metastases these lymph node metastases display increased resistance to the EGFR inhibitor erlotinib as compared to primary tumor NME cells when cultured under 3D organotypic conditions (fig. S1). Furthermore, the downstream EGFR signaling in these lymph node metastases also became dominated by STAT1 phosphorylation (fig. 1e). The diminution of EGFR through metastasis is further supported by previous reports from our lab and others showing EGFR downregulation in the RAS-transformed MCF10A progression series ^6,13,34^ To examine EGF-induced downstream signaling in this additional isogenic model of metastatic progression, we engineered the metastatic Ca1a cells to re-express EGFR using stable or doxycycline-inducible expression systems (fig. 1f and S2). EGF stimulation of Ca1a cells stably or transiently replenished with EGFR expression led to robust phosphorylation of EGFR and STAT1 and induction of apoptosis (fig. 1f, S2). Overall, these data indicate that enhanced STAT1 signaling downstream of EGFR activation correlates with EGF-mediated growth inhibition in metastatic BC.

### STAT1 is required for EGF-mediated apoptosis in metastatic BC

Previous studies indicate that STAT1 activation by EGF or other cytokines inhibits proliferation and induces apoptosis ^16–18^. Therefore, we sought to define the functional role of STAT1 downstream of EGFR activation in metastatic breast cancer cells. Indeed, EGF stimulation of both cell lines derived from lymph node metastases, Lym1 and Lym2, resulted in cell rounding and enhanced apoptosis as determined via a caspase 3/7 activity assay (fig. 2a and 2b). Depletion of STAT1 expression in the metastatic Lym1 cells using two different shRNA sequences prevented that ability of EGF to induce apoptosis (fig. 2c and 2d). Identical results were observed when STAT1 was depleted in the Lym2 cell line (data not shown). In these analyses, we also pharmacologically blocked Erk1/2 signaling through addition of trametinib While addition of trametinib alone had no effect on apoptosis, it did potentiate the ability of EGF to induce apoptosis in these cells (fig. 2d). Importantly, this effect was similarly dependent on STAT1 expression (fig. 2d) These data clearly implicate the functional involvement of STAT1 in EGF-induced apoptosis in metastatic BC.

**Fig. 2.**
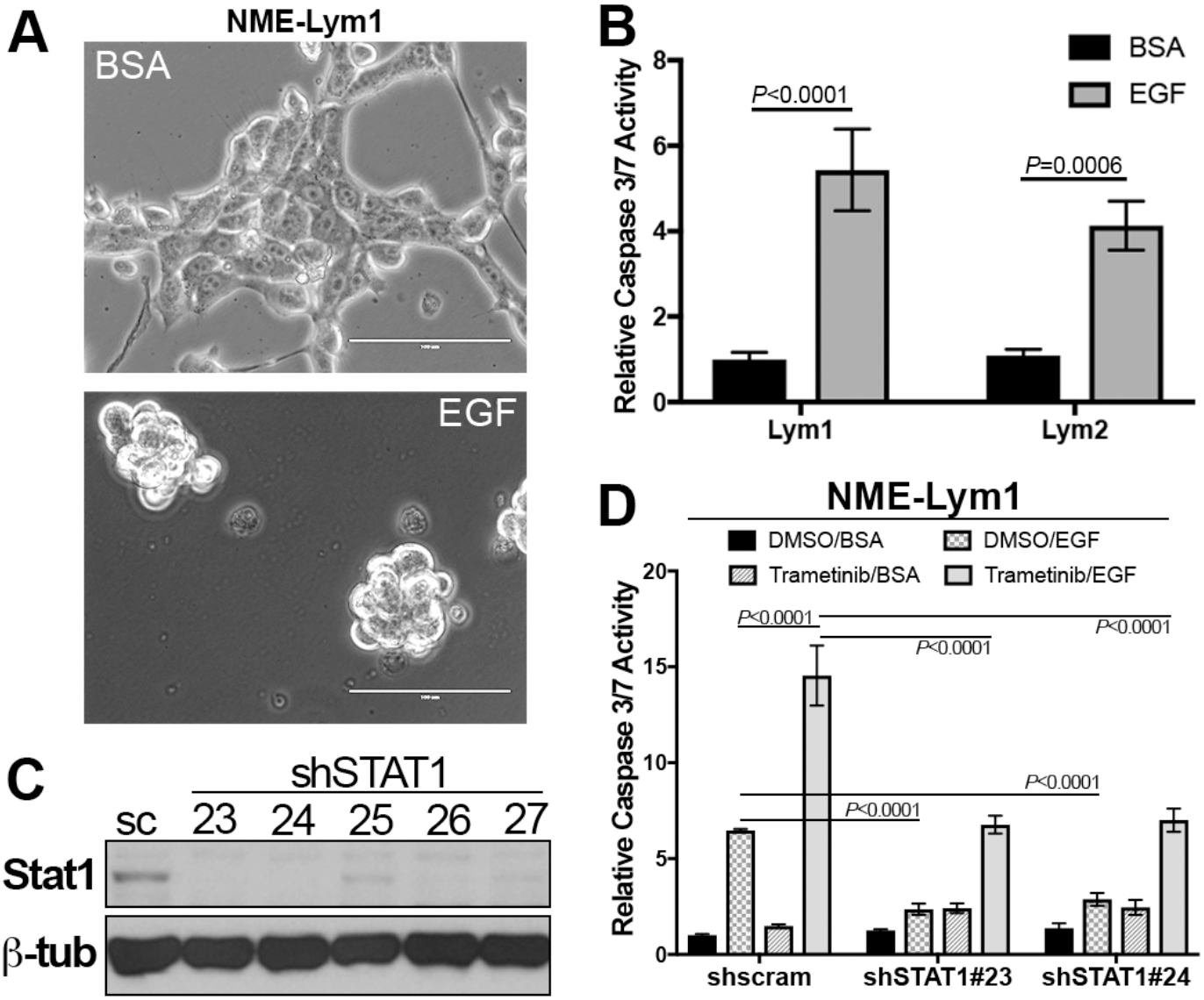
STAT1 is required for EGF-induced apoptosis in metastatic breast cancer cells (**A**) Lymph node metastases (NME-Lym1 and NME-Lym2) were stimulated with EGF (100 ng/ml) for 36 hours and imaged via phase contrast microscopy. (**B**) Following EGF stimulation as described in panel A cells were assayed for caspase 3/7 activity. (**C**) NME-Lym1 cells were constructed to stably express a scrambled control (sc) shRNA or various shRNAs (23-27) targeting STAT1. These cells were analyzed for STAT1 expression by immunoblot. Expression of b-tubulin (b-tub) served as a loading control. (**D**) Contorl (shscram) and STAT1 depleted (shSTAT1#23 and shSTAT1#24) NME-Lym1 cells were treated with EGF as described above and caspase 3/7 activity was assessed. Separate groups of cells were treated with the MEK1/2 inhibitor trametinib (100 nM) in the presence or absence of exogenous EGF (100 ng/ml) and these cells were similarly assayed for caspase 3/7 activity. Data in panels B and C are the mean ° SE of three independent experiments completed in triplicate resulting in the indicated *P* values.

### Nuclear STAT1 is accessed by EGFR through endocytosis

Our recent studies demonstrate an enhanced localization of EGFR in the nucleus of metastatic breast cancer cells as compared to primary tumor cells ^35^. Given the role of STAT1 in facilitating EGF-induced apoptosis in metastatic cells we next sought to investigate the subcellular localization of STAT1 under nonstimulated and EGF-stimulated conditions. Surprisingly, STAT1 was already localized to the nucleus in NME cells even before ligand stimulation (fig. 3a). Unfortunately, immunostaining these cells with a phospho-specific STAT1 antibody is not possible due to cross reactivity with phospho-EGFR epitopes. However, our whole-cell immunoblot analyses from Figure 1 indicate that prior to EGF stimulation, STAT1 is not phosphorylated (fig. 1b). Therefore, these data are consistent with previous reports that indicate a pool of STAT1 can exist in the nucleus in an unphosphorylated state ^36^. In contrast, ERK1/2 is primarily localized in the cytoplasm and only moves into the nucleus upon EGF stimulation, an event that is stabilized upon addition of leptomycin B to prevent nuclear export (fig. 3b). These data suggest that EGFR must gain access to the nucleus to phosphorylate STAT1. Indeed, EGFR internalization in endocytic vesicles can clearly be visualized upon EGF stimulation (fig. 3a). Moreover, closer examination of EGFR localization using super resolution microscopy revealed that in certain areas of the cell the plasma membrane is in direct physical contact with the nucleus. Therefore, we hypothesized that upon ligand-induced endocytosis from the plasma membrane a subpopulation of EGFR molecules has immediate access to the nuclear compartment (fig. 4a). To this end, we utilized an EGFR construct that contains alanine substitutions in the juxtamembrane dilysine motif (679-680-LL converted to AA). This construct is established to be signaling proficient from the plasma membrane, but deficient in endocytosis upon ligand engagement ^37^ Accordingly, the EGFR-AA construct was not able to induce phosphorylation of STAT1 in response to EGF (fig. 4b). Taken together these data consistently indicate that a subset of EGF receptors are able to translocate to the nucleus via endocytosis to phosphorylate STAT1.

**Fig. 3.**
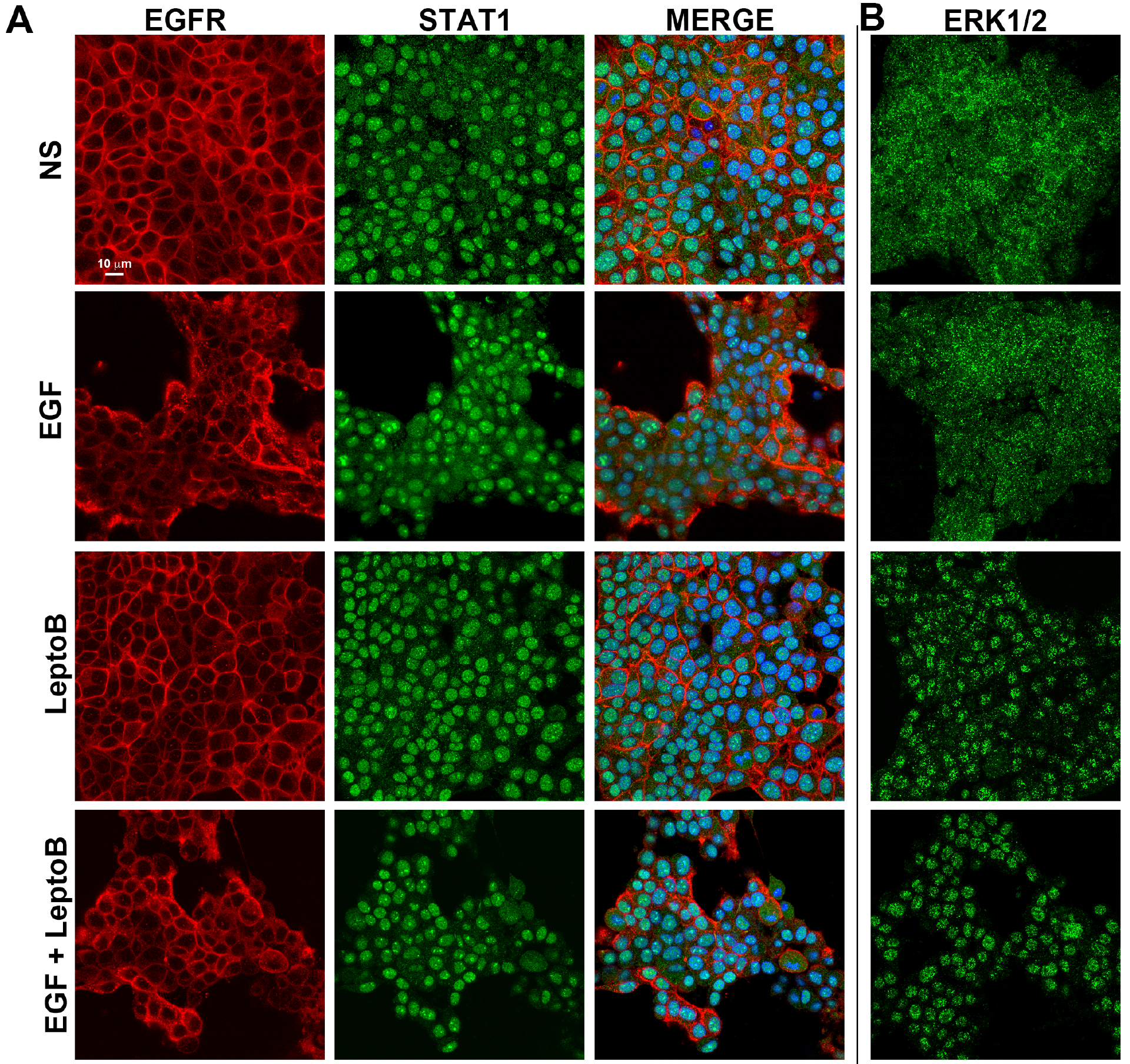
STAT1 is localized in the nucleus prior to phosphorylation. (**A**) NME cells were stimulated with EGF for 30 minutes in the presence or absence of the nuclear export inhibitor leptomycin B (LeptoB). These cells were subsequently analyzed by dual immunofluorescence and imaged via confocal microscopy for localization of EGFR and STAT1. These cells were counterstained with DAPI to visualize the nuclei. (**B**) Separate sets of cells were stimulated as in panel A and analyzed for localization of ERK1/2. Data in panels A and B are representative images from at least 10 fields of view over two independent experiments.

**Fig. 4.**
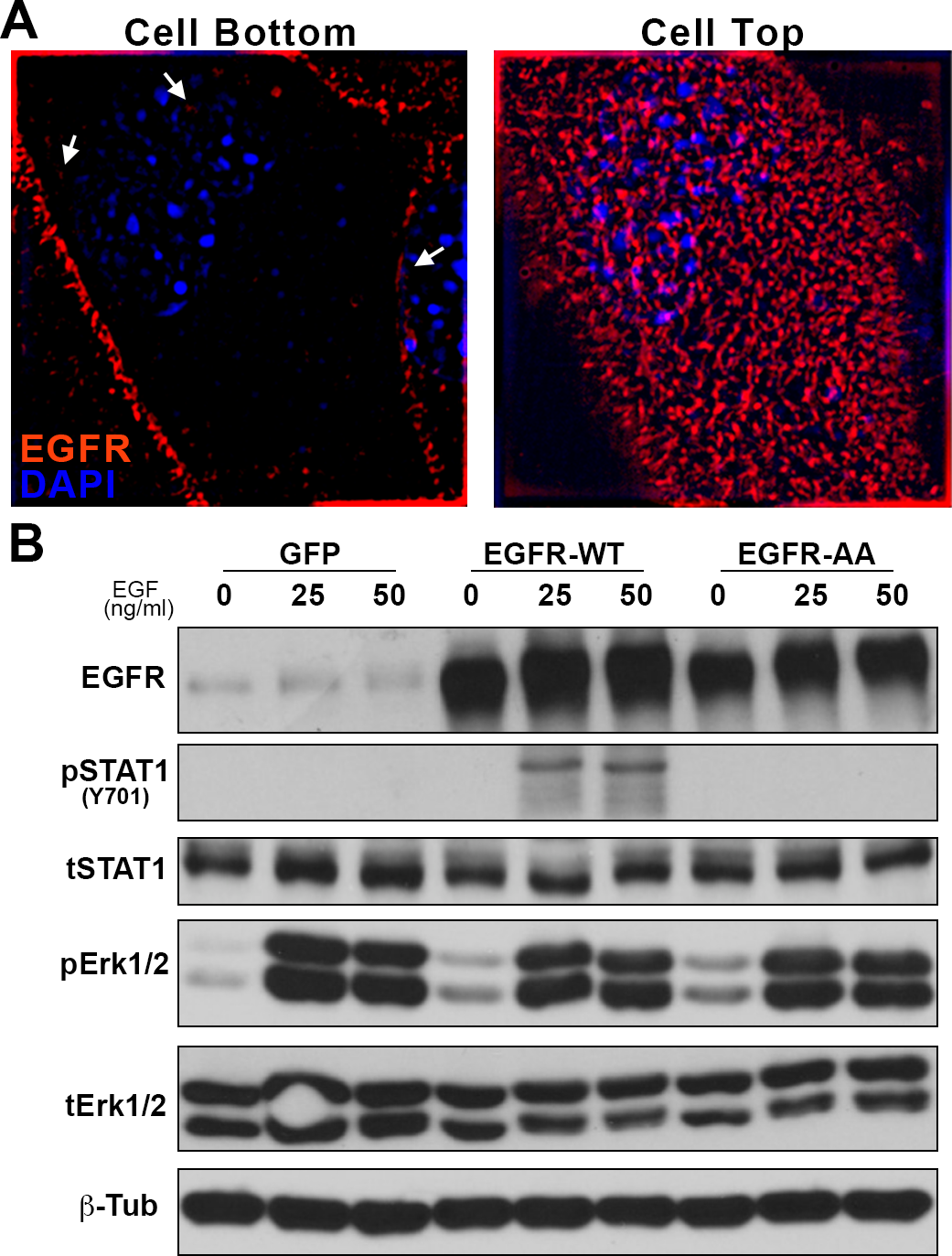
Endocytosis of EGFR is required for phosphorylation of nuclear STAT1. (**A**) NME cells were analyzed by immunofluorescence and imaged via super-resolution microscopy for localization of EGFR (100x objective). These cells were counterstained with DAPI to visualize DNA. Images of the same cell are shown for sections taken at interface of the cell with the coverslip (cell bottom) and top of the cell (cell top). Arrows indicate areas where EGF receptors appear to be in direct contact with the nucleus. (**B**) NMuMG cells expressing wild type EGFR (EGFR-WT), the endocytosis-deficient 679-680-AA mutant form of EGFR (EGFR-AA), or GFP as a control, were stimulated with EGF (50 ng/ml) for 30 minutes and analyzed for phosphorylation of STAT1 and ERK1/2. Analysis of total levels of EGFR, STAT1, ERK1/2, and β-tubulin (β-Tub) served as loading controls. Immunoblots are representative of at least three independent experiments.

### Pharmacological biasing of EGFR signaling promotes EGF-induced apoptosis in metastatic breast cancer cells

To specifically target the function of the subpopulation of EGF receptors that reach the nuclear compartment, we utilized our recently reported novel EGFR inhibitor ^35^. This chemical construct contains the EGFR tyrosine kinase inhibitor gefitinib linked to a peptoid moiety encoding the SV40 nuclear localization sequence (NLS-gefitinib). This approach leads to robust accumulation of geftinib in the nucleus ^35^. Consistent with the notion that only endocytosed EGFR molecules have access to the nuclear pool of STAT1, pretreatment of NME-LM1 cells with this chimeric NLS-gefitinib molecule led to potent blockade of EGF-induced STAT1 phosphorylation without affecting phosphorylation of ERK1/2 (fig. 5a). In contrast, pretreatment with trametinib drastically alters the STAT1:ERK1/2 activation ratio by completely preventing downstream phosphorylation of ERK1/2, while leading to a slight increase in EGF-induced phosphorylation of STAT1 (fig. 5a).

**Fig. 5.**
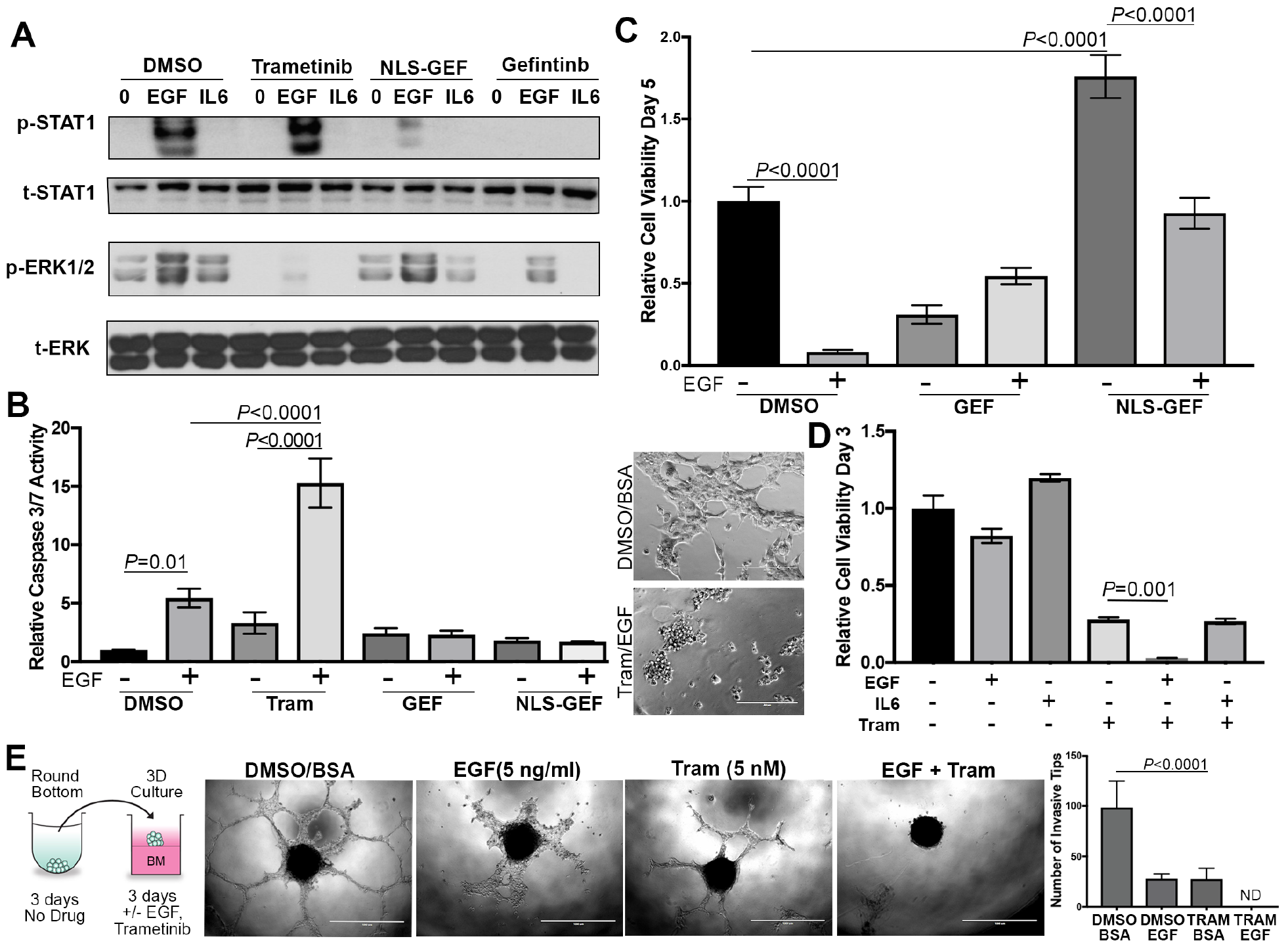
Inhibition of ERK1/2 signaling augments EGF-induced apoptosis in metastatic breast cancer cells. (**A**) Lung metastatic (NME-LM1) cells were pretreated with trametinib, gefitinib, or a nuclear localization sequence-gefitinib conjugate (NLS-GEF) and then stimulated with EGF (50 ng/ml) for 30 minutes. These cells were subsequently analyzed for phosphorylation of STAT1 and ERK1/2. IL6 and BSA (0) served as protein stimulation controls and total levels of STAT1 and ERK1/2 were assessed as loading controls. (**B**) NME-LM1 cells were stimulated with EGF (100 ng/ml) in the presence or absence of trametinib (Tram), gefitinib (GEF), or the nuclear localization sequence-gefitinib conjugate (NLS-GEF). Following 24 hours of treatment, these cells were assayed for caspase 3/7 activity. (Inset) Representative images of cells under control and Tram/EGF stimulation are shown. (**C**) As in panel B NME-LM1 cells were stimulated with EGF (50 ng/ml) in the presence of the indicated inhibitors for 5 days at which point changes in cellular viability were quantified. (**D**) NME-LM1 cells were stimulated as indicated for 3 days at which point cell viability was quantified. (**E**) NME-LM1 tumorspheres were formed in round bottom wells and subsequently transferred to a hydrogel layer of basement membrane in the presence or absence of EGF and trametinib. Tips of invading cellular branches were quantified. Data in panels B-E are the mean ±SE of three independent experiments completed in triplicate.

To quantify the biological implications of these events, we again analyzed caspase 3/7 activation following EGF treatment in the presence or absence of these inhibitors (fig. 5b). Consistent with our findings in figure 2 implicating the role of STAT1 in facilitating EGF-induced apoptosis, addition of gefitinib-NLS blocked EGF-induced activation of caspase 3/7 (fig. 5b). As we observed with lymph node metastases, cotreatment of these lung metastases with trametinib led to a robust increase in EGF-induced caspase 3/7 activity (fig. 5b). Also, NLS-gefitinib was capable of preventing EGF-mediated growth inhibition in the NME-LM1 cells, and NLS-gefitinib increased cell viability under nonstimulated conditions (fig. 5c). In contrast, addition of EGF augmented the growth suppressive effects of trametinib in a shorter-term assay (fig. 5d). Consistent with our shRNA depletion studies, the ability of EGF to induce apoptosis is likely dependent on STAT1 since addition of IL-6, a specific activator of STAT3, did not alter trametinib-induced growth inhibition (fig. 5d). All of these events could be replicated using the LM2 metastatic variant (fig. S3). The only noted difference in the LM2 cells was that no caspase 3/7 activity could be quantified with EGF alone, but again addition of EGF and trametinib led to significantly increased apoptosis as compared to trametinib alone (fig. S3a)

As cells escape primary mammary tumors, a particularly aggressive subpopulation is able to survive the non-adherent conditions in blood/lymphatic circulation, and these cells are ultimately responsible for colonizing vital organs. To recapitulate these events in an *in vitro* assay, we generated spheroids of metastatic cells in non-adherent conditions and then transfered these spheroids onto a bed of reconstituted basement membrane (fig. 5e). Using this approach, the metastatic NME-LM1 tumorspheres form highly invasive, multicellular branches over a period of three days (fig. 5e). Addition of physiological amounts of EGF or nanomolar concentrations of trametinib partially blocked these events, but a combination of the growth factor and trametinib completely prevented invasive growth of these highly metastatic cells. Therefore, biasing EGFR signaling toward STAT1 using downstream inhibitors can enhance the apoptotic potential of this pathway.

### Pharmacological biasing of EGFR signaling fundamentally changes response to EGF in primary tumor cells

We next sought to determine if biasing EGFR signaling could change the EGF response of primary tumor cells from proliferative to apoptotic. Therefore, we treated the nonmetastatic NME cells with EGF in the presence of NLS-gefitinib, unconjugated gefitinib, or trametinib. As observed in figure 1, very little activation of STAT1 in NME cells occured upon EGF stimulation, however this pathway was greatly enhanced upon inhibition of MEK1/2 with trametinib (fig. 6a). Consistent with the notion that the ERK1/2:STAT1 activation ratio dictates the proliferative versus apoptotic outcome of EGF, we observed a drastic induction of apoptosis in the NME cells upon co-adminstration of EGF and trametinib, whereas either treatment alone did not produce any caspase activation (fig. 6b and 6c). Moreover, these effects could not be produced in control cells expressing normal amounts of EGFR, or in cells expressing the EGFR-AA variant that is deficient in STAT1 activation (fig. S4 and fig. 6c). Finally, using our 3D spheroid assay described in figure 5 we could illustrate the non-metastatic nature of the NME cells as they fail to form any invasive structures (fig. 6d). However, we could quantify an increase in spheroid size with addition of exogenous EGF, a result that was completely prevented upon addition of trametinib (fig. 6d and 6e). Moreover, co-treatment with EGF and trametinib lead to the appearance of apoptotic bodies around the tumorspheres (fig 6d; inset). These data indicate that biasing signaling away from ERK1/2 can fundamentally alter how cells respond to ligand-mediated activation of EGFR.

**Fig. 6.**
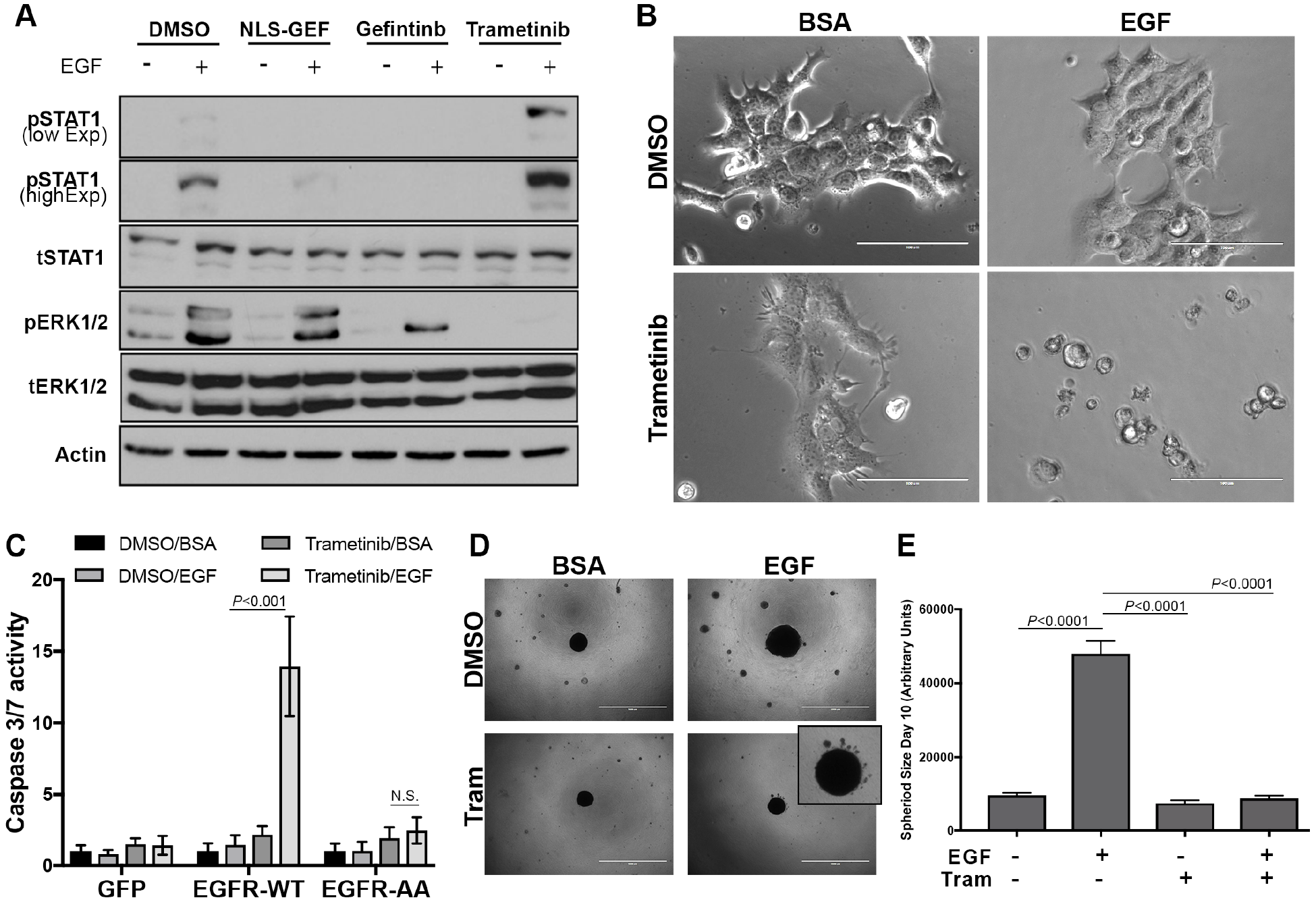
Inhibition of Erk1/2 signaling fundamentally changes response to EGF in primary tumor cells. (**A**) NME cells were stimulated with EGF (50 ng/ml) for 30 minutes in the presence or absence of trametinib, gefitinib or the gefitinib nuclear localization sequence-conjugate (NLS-GEF). These cells were then analyzed for phosphorylation of STAT1 and ERK1/2. Expression of total STAT1, ERK1/2 and Actin served as loading controls. (**B**) Representative brightfield photomicrographs of NME cells stimulated for 24 hours with EGF (50 ng/ml), trametinib (5 nM) or the combination. (**C**) Control (GFP) NMuMG cells or those cells expressing wild type EGFR (EGFR-WT) or the 679-680-AA variant of EGFR (EGFR-AA) were stimulated with EGF in the presence or absence of trametinib as in panel B and assessed for Caspase 3/7 activity. (**D**) Representative photomicrographs of NME tumorspheres cultured under 3D hydrogel conditions in the presence or absence of exogenous EGF (5 ng/ml) and trametinib (5 nM). (**E**) Quantification of NME spheroid size under the conditions described in panel D. Data in panels C and E are the mean ± SE of three independent experiments completed in triplicate.

### EGFR:STAT1 signaling augments trametinib-induced growth inhibition

To extend our findings to other breast cancer models of EGFR signaling we applied a similar EGF-trametinib treatment combination to our dox-inducible model of EGFR expression in the metastatic Ca1a cells. When EGFR expression was induced with dox we observed STAT1 phosphorylation in response to EGF stimulation (fig. 7a). Consistent with the Ca1a cells being transformed by a constitutively active form of RAS, we did not observe any further ERK1/2 phosphorylation upon EGF stimulation (fig. 7a). However, EGF-induced phosphorylation of STAT1 was enhanced in the presence of trametinib concentrations capable of completely blocking this constitutive phosphorylation of ERK1/2 (fig. 7a). Using these cells, we observed a dose dependent induction of EGFR expression with dox (fig. 7b). This transient induction resulted in an EGFR-dependent inhibition of cell growth in the presence of EGF and trametinib that was not observed in the presence of trametinib alone (fig. 7c). We similarly assessed the MDA-MB-468 (468) model of triple negative breast cancer. As we observed in our metastatic cells and consistent with previous reports, treatment of the 468 cells with EGF induced phosphorylation of STAT1 (fig. 7d; ^17^). EGF-induced phosphorylation of STAT1 was again completely blocked by our nuclear localized EGFR inhibitor, a condition that has no effect on ERK1/2 phosphorylation (fig. 7d). Importantly, pretreatment with trametinib at a concentration that completely prevented ERK1/2 phosphorylation enhanced EGF-induced STAT1 signaling (fig. 7d). Finally, combined treatment with EGF and trametinib lead to a potent inhibition of cell viability as compared to either treatment alone (fig. 7e). Such combination effect was not observed with IL6 which has been shown to inhibit the growth of 468 cells by conferring stemlike properties but is unable to induced STAT1 phosphorylation.

**Fig. 7.**
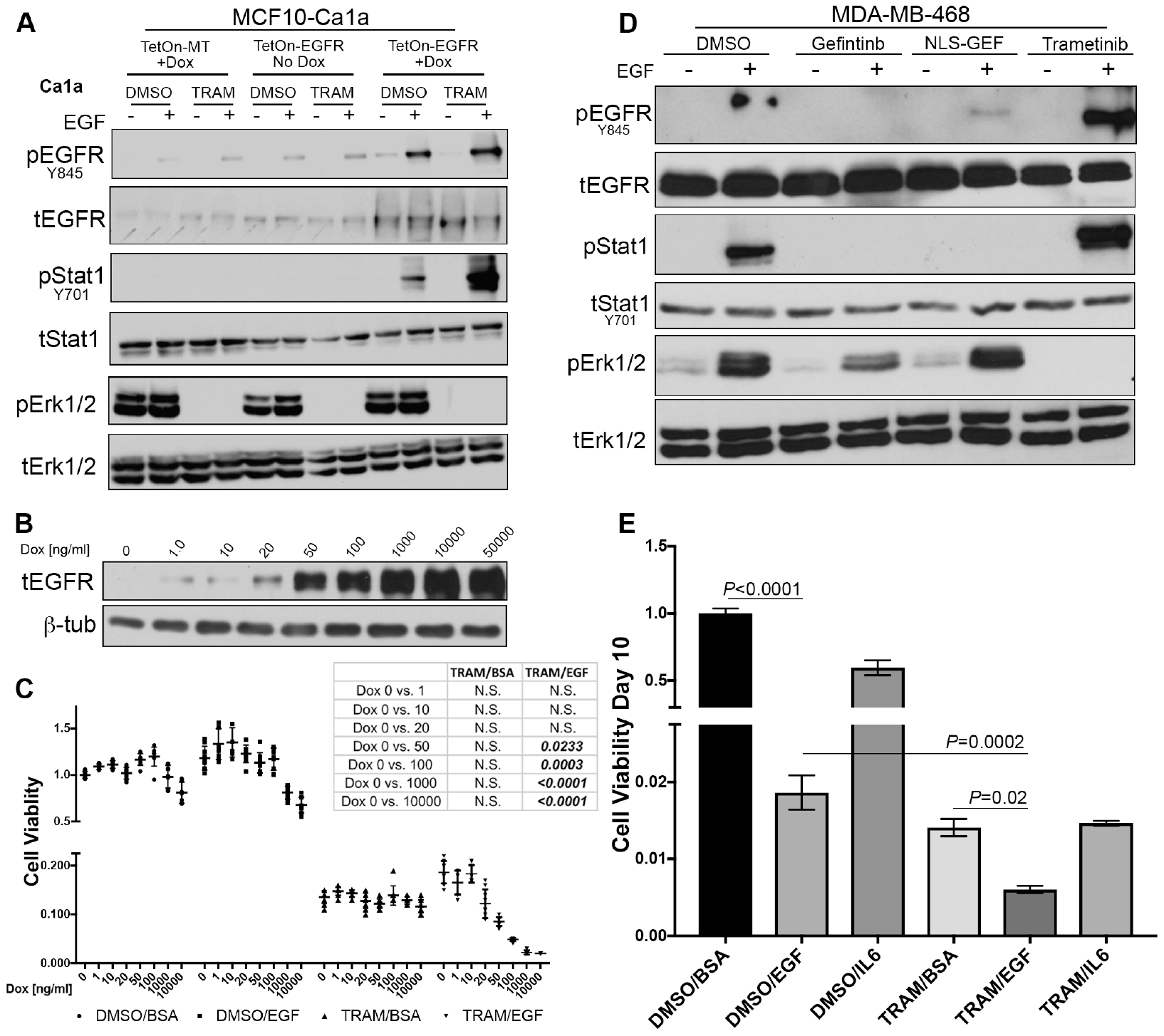
EGF enhances the growth inhibitory effect of trametinib. (**A**) Metastatic MCF10-Ca1a cells were constructed to express a control (TetOn-MT) or EGFR (TetOn-EGFR) encoding vector under the control of doxycycline (Dox) inducible promoter. These cells were pretreated with Dox for 24 hours and subsequently stimulated with EGF (50 ng/ml) for 30 minutes in the presence or absence of trametinib (TRAM). Phosphorylation of EGFR, STAT1 and ERK1/2 was assessed. Expression of total EGFR, STAT1 and ERK1/2 served as loading controls. (**B**) MCF10-Ca1a TetOn-EGFR cells were stimulated with the indicated concentrations of Dox for 24 hours and assessed for expression of EGFR. β-Tub served as a loading control. (**C**) MCF10-Ca1a TetOn-EGFR cells were stimulated with the indicated concentrations of Dox for 10 days in the presence or absence of EGF and trametinib and cell viability was quantified. *Chart inset:* The resultant *P* values of ANOVA analyses comparing the indicated treatment groups under control (Dox 0) and Dox conditions. (**D**) The MDA-MB-468 cells were stimulated with EGF (50 ng/ml) for 30 minutes in the presence or absence of trametinib and assessed for phosphorylation of EGFR, STAT1 and ERK1/2. Expression of total EGFR, STAT1 and ERK1/2 served as loading controls. (**E**) The MDA-MB-468 cells were grown for a period of 10 days in the presence or absence of EGF (50 ng/ml), IL6 (20 ng/ml), trametinib (5 nM) of the indicated combinations at which point cell viability was quantified. The indicated groups were analyzed by T-test resulting in the indicated *P* values. Data in panels A, B and D are representative of three independent analyses, and data in panels C and E are the mean ± SE for three independent experiments completed in triplicate.

## Discussion

EGFR activation is upstream of multiple signal transduction pathways. The differential activation of particular pathways in response to ligand leads to oncogenic versus apoptotic signals in specific cell types ^5,38,39^ We have previously reported that EGFR function paradoxically changes from oncogenic to apoptotic after *in vivo* metastasis of BC cells ^19^. In the current study, we demonstrate that ligand-mediated EGFR activation ultimately results in a nuclear STAT1-dependent apoptosis of metastatic BC cells. This fundamental change in response to EGF through breast cancer progression led us to address the hypothesis that pharmacological biasing of downstream signaling could reveal apoptotic EGFR signaling, even in early stage breast cancer. This concept is supported by recent studies in the fields of G-protein coupled receptor signaling and receptor tyrosine kinase signaling which indicate that differential ligand stimulation leads to biased downstream pathway activation and therefore unique biological outputs ^40,41^. These previous studies have focused on unnatural ligands and allosteric modulation of receptors. In contrast, our work herein demonstrates that unique cellular outcomes in response to an endogenous ligand can be manifested when specific downstream pathways are pharmacologically interdicted.

Recent studies indicate that constitutive EGFR signaling induced upon receptor mutations are distinct and mutually exclusive as compared to ligand-induced signaling ^42^. Our findings indicate that direct pharmacological targeting of WT-EGFR using our nuclear localized gefitinib conjugate completely prevents STAT1 activation, leading to enhanced cell growth as this compound has no effect on the ability of EGFR to signal to ERK1/2. Furthermore, in several instances we observed unconjugated gefitinib prevents STAT1-mediated apoptotic signaling downstream of ligand activated WT-EGFR at a much lower concentration than is required for inhibition of ERK1/2-mediated proliferative signaling. These data suggest that complete to near-complete blockade of EGFR function is required before an anti-tumorigenic response would be expected in breast cancer cells bearing high levels of WT-EGFR undergoing ligand-mediated signaling. Indeed, complete pharmacological blockade of a target molecule is challenging if not impossible to achieve *in vivo.* This concept that incomplete inhibition of WT-EGFR is biasing signaling toward proliferative ERK1/2 signaling is completely consistent with clinical failure of EGFR inhibitors in metastatic breast cancer ^20^.

Our recent studies demonstrate that metastatic cells increase their nuclear pool of EGFR ^35^. These data are consistent with the findings here demonstrating an enhanced ability of metastatic cells to access and phosphorylate nuclear STAT1 in response to EGF stimulation. The potential mechanisms by which metastatic cells increase their nuclear pool of EGFR are potentially numerous (42–44). However, recent data suggest that more migratory cells undergo constant nuclear rupture, and these cells repair these events by using components of endosomal sorting complexes ^46^. Together with our findings using the EGFR-AA construct, these data suggest a mechanism in which more migratory and metastatic breast cancer cells undergo an increased rate of nuclear rupture and repair and thus sample more activated EGFR molecules from endosomes. These events would lead to an increased pool of nuclear EGFR and enhanced interaction of these receptors with the nuclear pool of STAT1. Finally, several of the model systems interrogated herein demonstrate that trametinib enhances EGFR:STAT1 signaling. Our recent studies indicate that EGFR signaling is regulated in metastatic cells via expression of the EGFR inhibitory molecule Mig6 ^19^. Expression of Mig6 is driven via ERK1/2, constituting a physiologic negative feedback on EGFR activation ^41,48^. Although not evaluated here, trametinib may serve to short circuit this negative feedback by decreasing Mig6 and allowing unabated activation of alternate, apoptotic signaling downstream EGFR such as STAT1.

In conclusion, our studies broadly illustrate the importance of understanding the cellular outcomes of cytoplasmic kinase inhibitors not only in terms of the pathway they are targeting, but also in terms of changes they insight to alternate signal transduction pathways induced from shared upstream receptors. These data support current clinical trials evaluating the efficacy of trametinib in the treatment of metastatic breast cancer (NCT02900664, NCT03065387). However, our results argue against the concurrent use of EGFR kinase inhibitors in these patients as this will block apoptotic, EGFR:STAT1 signaling, limiting the apoptotic effect of trametinib treatment. Current studies in the lab are exploring therapeutic approaches to enhance the antitumor effects of trametinib through specific augmentation of EGFR:STAT1 signaling.

## Acknowledgments

The authors thank the members of the Wendt lab for critical reading of the manuscript. This research was supported in part by National Institutes of Health (R01CA207751) to MKW, the American Cancer Society (RSG-CSM130259) and METavivor foundation to MKW. Purdue Center for Cancer Research via an NIH NCI grant (P30CA023168). The authors also acknowledge the use of the facilities of the Bindley Bioscience Center, a core facility of the NIH-funded Indiana Clinical and Translational Sciences Institute.

---

**fig. S1.**
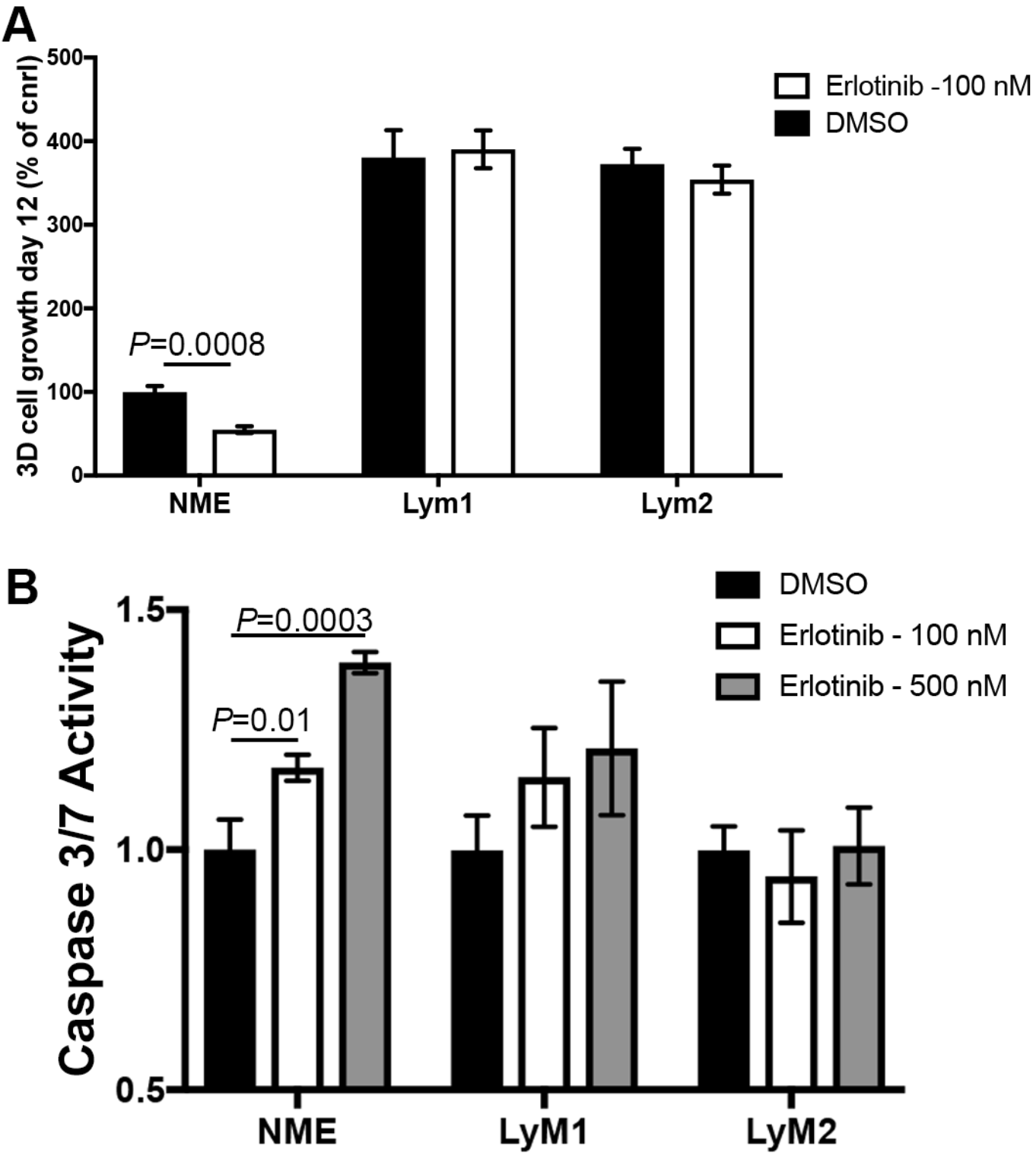
EGFR-transformed lymph node metastases are resistant to inhibition of EGFR kinase activity. (**A**) EGFR transformed mammary epithelial cells (NME) and their isogeneic lymphnode-derived metastatic counterparts (Lym1 and Lym2) expressing firefly luciferase were cultured under 3D conditions in the presence or absence of the EGFR inhibitor erlotinib for 12 days at which point cellular viability was quantified by bioluminescence. (**B**) The cells described in panel A were cultured on 2D tissue plastic and treated for 24 hours with the indicated concentrations of erlotinib and subsequently analyzed for caspase 3/7 activity. Data are the mean ± SE of three separate experiments completed in triplicate, resulting in the indicated *P* values.

**fig. S2.**
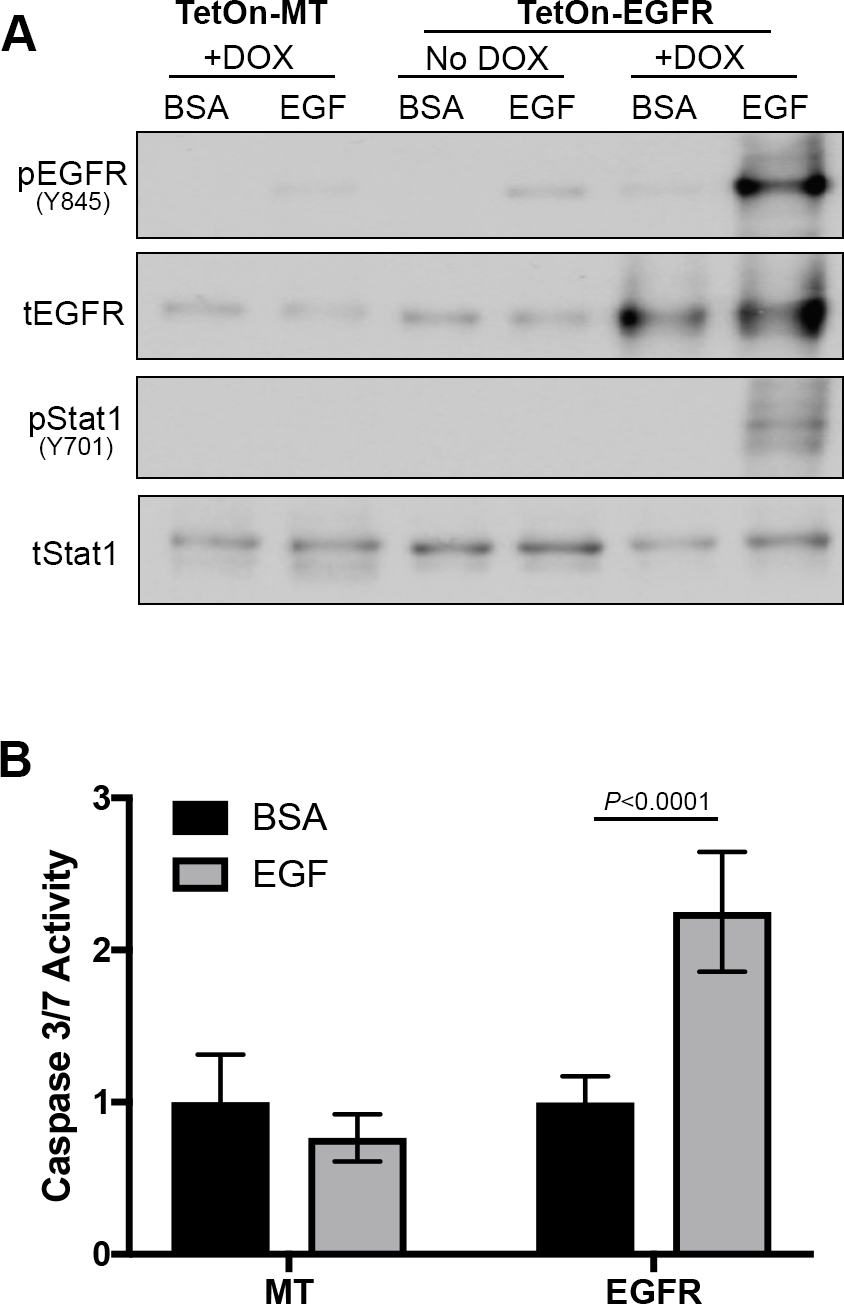
Induction of EGFR in metastatic breast cancer cells is sufficient for ligand induced STAT1 phosphorylation and apoptosis. (**A**) Metastatic MCF10-Ca1a cells were constructed to express EGFR under the control of a tetracycline-induced promoter. Following a 24-hour induction with doxycycline (DOX; 1 αg/ml) these cells were stimulated with EGF (50 ng/ml) for 30 minutes and assayed for phosphorylation of EGFR and STAT1. Expression of total EGFR and STAT1 served as loading controls. (**B**) MCF10-Ca1a cells were constructed to stably express EGFR. These cells were stimulated with EGF (50 ng/ml) for 24 hours and assayed for caspase 3/7 activity.

**fig. S3.**
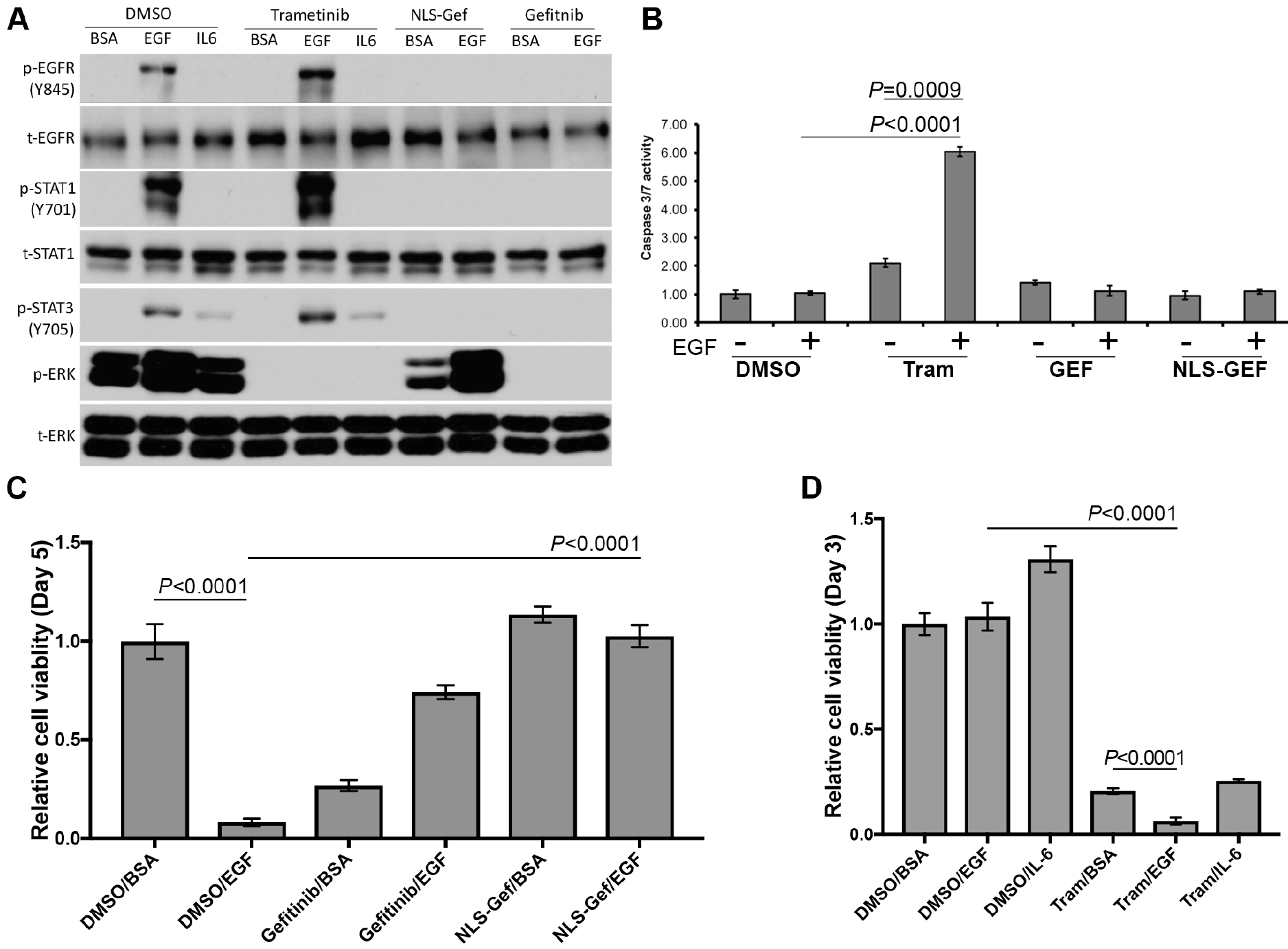
Inhibition of ERK1/2 signaling causes EGF-induced apoptosis in lung metastatic breast cancer cells. (**A**) Lung metastatic (NME-LM2) cells were pretreated with trametinib, gefitinib, or a nuclear localization sequence-gefitinib conjugate (NLS-GEF) and then stimulated with EGF (50 ng/ml) for 30 minutes. These cells were subsequently analyzed for phosphorylation of EGFR, STAT3, STAT1 and ERK1/2. BSA (0) served as a protein stimulation control and total levels of EGFR, STAT1 and ERK1/2 were assessed as loading controls. (**B**) NME-LM2 cells were stimulated with EGF (100 ng/ml) in the presence or absence of the indicated inhibitors (Tram = trametinib, GEF = gefitinib, NLS-GEF = gefitinib conjugated to a nuclear localization sequence). Twenty-four hours later cells were assessed for caspase 3/7 activity. (**B**) LM2 cells were treated with trametinib (Tram; 5 nM) in the presence or absence of EGF (50 ng/ml) or IL6 for three days at which point cell viability was quantified. (**C**) LM2 cells were stimulated with EGF (50 ng/ml) in the presence or absence of the indicated inhibitors for 5 days at which point cell viability was quantified. All data are the mean ± SE of three separate experiments completed in triplicate resulting the indicated *P* values.

**fig. S4.**
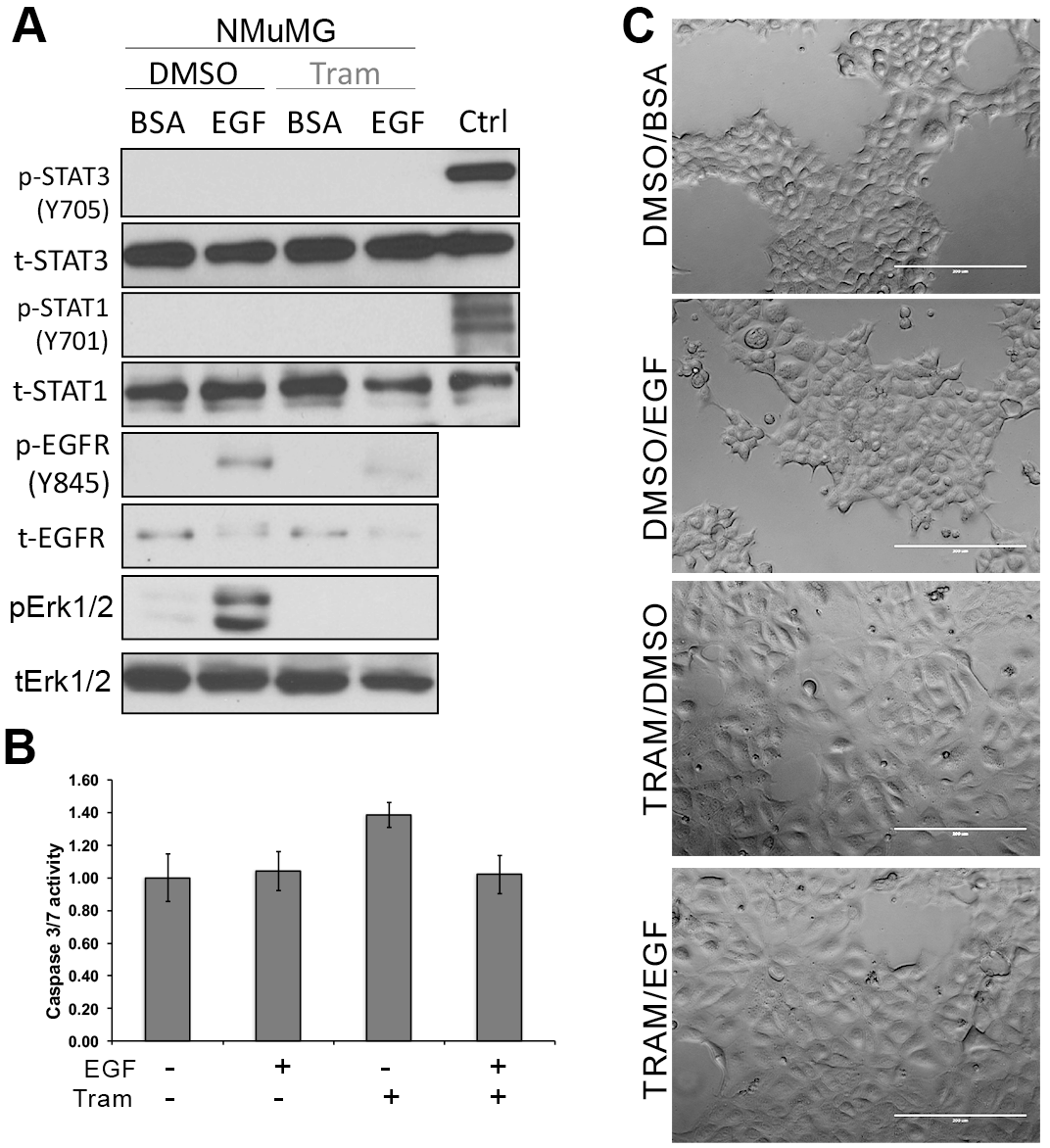
A combination of EGF and trametinib does not induce STAT1 phosphorylation or apoptosis in normal mammary epithelial cells. (**A**) Nontransformed NMuMG cells were stimulated with EGF (50 ng/ml) for 30 minutes in the presence or absence of trametinib (tram; 100 nM). These cells were lysed and analyzed for phosphorylation of STAT1, ERK1/2 and EGFR. Expression of total EGFR, STAT1 and ERK1/2 served as loading controls. (**B**) NMuMG cells were stimulated with EGF (100 ng/ml) in the presence and absence of trametinib (Tram; 100 nM) for 24 hours and these cells were assessed for caspase 3/7 activity. Data are the mean ± SE of three separate experiments completed in triplicate. (**C**) Phase contrast photomicrographs of NMuMG cells treated as described in panel B.

